# Mouse embryonic stem cells can differentiate via multiple paths to the same state

**DOI:** 10.1101/124594

**Authors:** James A. Briggs, Victor C. Li, Seungkyu Lee, Clifford J. Woolf, Allon M. Klein, Marc W. Kirschner

## Abstract

In embryonic development, cells must differentiate through stereotypical sequences of intermediate states to generate mature states of a particular fate. By contrast, direct programming can generate similar fates through alternative routes, by directly expressing terminal transcription factors. Yet the cell state transitions defining these new routes are unclear. We applied single-cell RNA sequencing to compare two mouse motor neuron differentiation protocols: a standard protocol approximating the embryonic lineage, and a direct programming method. Both undergo similar early neural commitment. Then, rather than transitioning through spinal intermediates like the standard protocol, the direct programming path diverges into a novel transitional state. This state has specific and abnormal gene expression. It opens a ‘loop’ or ‘worm hole’ in gene expression that converges separately onto the final motor neuron state of the standard path. Despite their different developmental histories, motor neurons from both protocols structurally, functionally, and transcriptionally resemble motor neurons from embryos.

## Introduction

Embryonic development proceeds through defined intermediate states, such as germ layer intermediates, and lineage-specific progenitors. Intermediates bifurcate into multiple states over time, and specialize their behaviors, ultimately producing a lineage tree that defines each mature cell type by a particular sequence of intermediates. This was first appreciated through classical lineage tracing and cell ablation studies. These studies showed that specifically labeled intermediate states generate stereotyped sets of downstream cell types, and that these downstream cell types fail to form if an intermediate that is upstream in their lineage is ablated^1,2^. Furthermore, embryos in general do not produce a mature cell type through multiple differentiation paths.

In contrast to this rigid and hierarchical process, recent protocols that experimentally directly program cell fate suggest that the exact sequence of intermediates defining a lineage may be more flexible^3-11^. These studies reveal that mature cell states can be reached through paths that do not involve activation of the intermediate progenitor genes that are essential in embryos. Mouse embryonic stem cells (mESCs), for example, can be converted into motor neurons (MNs) by a process that involves overexpression of three transcription factors, Ngn2+Isl1+Lhx3^3,12^, and that never expresses the neural progenitor transcription factors Sox1 and Olig2^3^. mESCs can also be driven rapidly into a terminal muscle phenotype without normal upregulation of intermediate genes such as Pax7 and Myf5, through a combination of cell-cycle inhibition and MyoD overexpression^4^. This plasticity of differentiation extends further, to the interconversion of mature cell states. Fibroblasts can be converted into mature neuron phenotypes^6,11^, including MNs^5^, seemingly without completely dedifferentiating and retracing the embryonic lineage, as indicated by lack of expression of specific core pluripotency (Oct4, Sox2 and Nanog) and neural progenitor genes (Nestin)^5^.

Although these direct programming (DP) experiments imply the existence of differentiation paths that differ from those in embryos, much of what actually occurs in these new programs remains mysterious. Does DP bypass normal intermediates by short-circuiting the natural lineage, or does it transition through alternative intermediates (Fig. 1a)? Does it diverge only briefly to bypass specific early or late states, or does it utilize an entirely distinct path (Fig. 1b)? And can DP converge fully to the same final state that is produced in embryos despite taking an alternative path (Fig. 1c)? These questions have been challenging to answer in part due to the high degree of heterogeneity in direct programming experiments, where unbiased bulk measurements of global gene expression obscure changes, and also because marker genes allowing the isolation of new but potentially important DP-specific intermediates are not known in advance. Here we aimed to overcome these issues by applying single cell RNA sequencing to compare the gene expression trajectories of DP and growth factor guided differentiation of mESCs over time into MNs. Our core research questions are summarized in Figure 1.

**Fig. 1.**
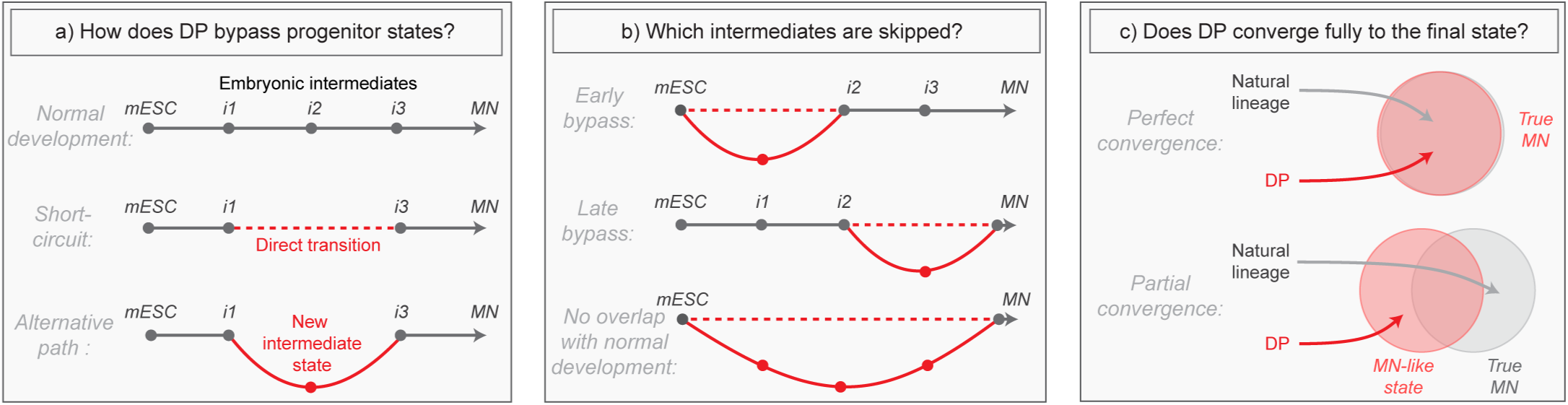
Summary of core research questions. a) Direct programming could in principle skip progenitor states in one of two ways. It may either transition from early states directly into later states through a ‘short-circuit’ of the natural lineage, or it might utilize an alternative path involving new and potentially abnormal or unstable intermediate states. In the schematic, a conceptual depiction of the natural lineage is shown first, followed by two lineages that bypass the *i2* intermediate from the normal lineage, either through short-ciruit or through a new alternative intermediate, b) DP might diverge from the natural lineage only briefly, or it might access an entirely distinct path. In the schematic, three different possibilities are shown: i) DP bypasses specific early intermediates, and then converges to the final state through a conserved seqeunce of terminal cell state transitions; ii) DP transitions through conserved early stages of differentiation, before diverging into an alternate path that converges separately to the final state; iii) DP takes an entirely distinct path with no resemblance to the natural lineage. Dashed lines represent a potential direct transition while solid lines represent an alternate path (as shown in the left panel), c) It is possible that the price paid for taking an alternate differentiation path during direct programming is that cells retain subtle gene expression defects, thus converging only partially to the desired final state. Alternatively, convergence could be near perfect which would imply that differentiation into mature cell states is at least partially history independent. In the schematic, circles are an abstract representation of gene expression state and are used to represent the extent of overlap between MNs generated by either DP or by the natural lineage.

## Results

### Dissection of two MN differentiation protocols using InDrops single cell RNA sequencing

We compared two *in vitro* differentiation protocols that convert mESCs into MNs. Spinal MNs were chosen for study because these protocols have been highly optimized. The first, standard protocol (SP) is a widely used method that attempts to recapitulate the known embryonic intermediates through sequential exposure of developmental signals (Fgfs, Retinoic Acid, and Sonic hedgehog)(Fig. 1A)^13,14^. It provides an approximation of the lineage through which motor neurons develop in the embryo. The second, direct programming (DP) protocol involves driving the expression of transcription factors (Ngn2+Isl1+Lhx3)^3,12^ that characterize the mature motor neuron state and at the same time favoring and stabilizing the G1 state by incubating in a growth factor free medium^4^.

We used single-cell RNA sequencing (InDrops)^15^ to track the differentiation trajectories of both protocols over time (Figs. 1B and 1C). Single-cell data has emerged as a powerful way to trace differentiation processes, particularly in populations that are not pure and that contain rare intermediates^11,16-20^. We profiled a total of 4,590 cells sampled from early (day 4/5) and late (day 11/12) timepoints for each protocol, and also used our previously published data from 975 mES cells^15^. To visualize the single cell data and identify cell states we applied t-distributed stochastic neighbor embedding (tSNE) to reduce dimensionality^21,22^, defined cell states using an unsupervised density gradient clustering approach, and then found specific marker genes with known annotations to reveal the identity of each state (Fig. 2B - 2E; Supp. Figs. S1 and S2; Supplementary methods). For each protocol, the dominant feature was a continuous gene expression trajectory sweeping across the 2D plot. These trajectories correlate with chronology: they begin with mESCs, pass through neural progenitor states and terminate in mature MN states. In both protocols we also observed a mixture of off-target differentiation byproducts from all three germ layers.

**Fig. 2.**
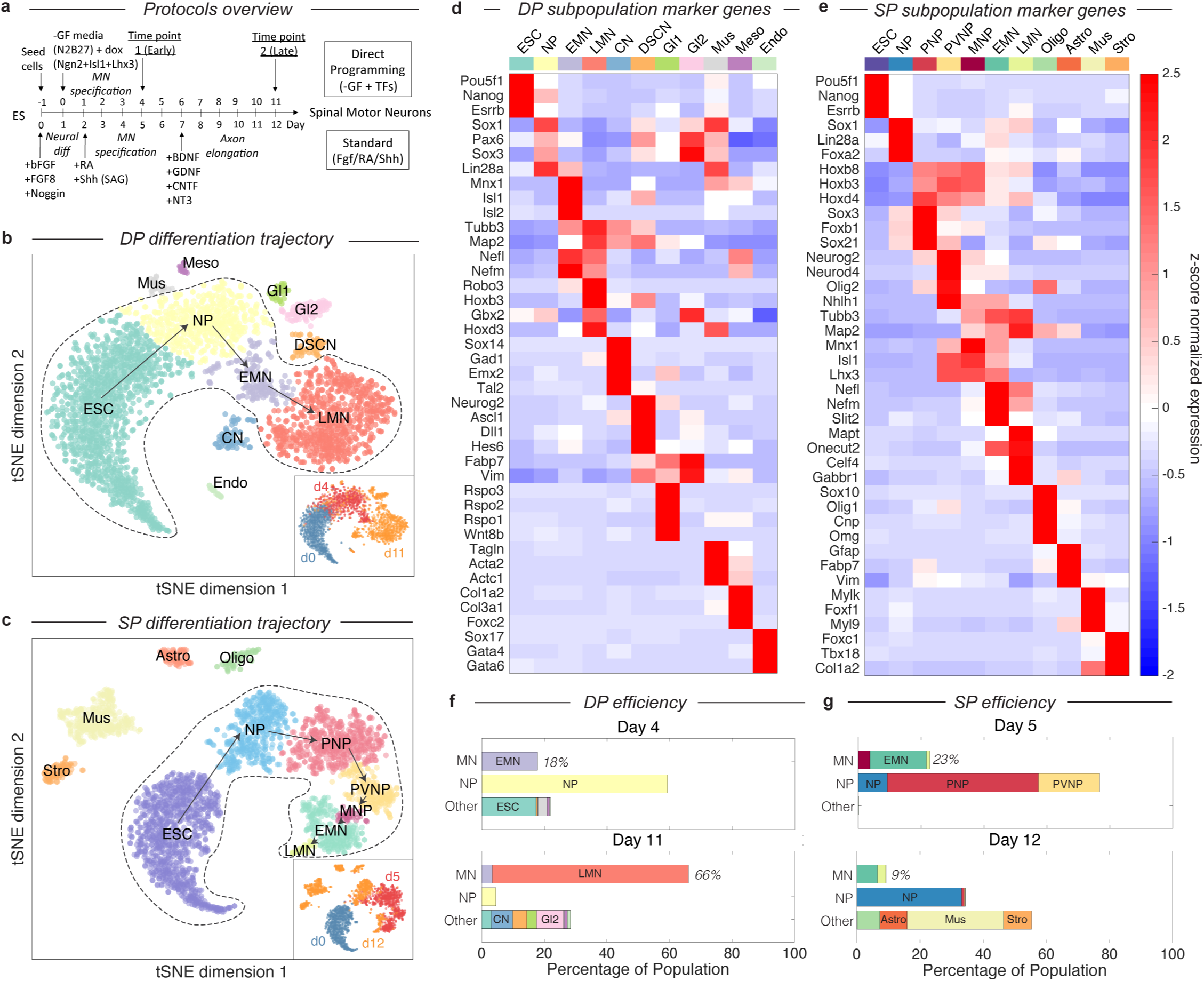
Dissection of DP and SP motor neuron differentiation strategies using InDrops single cell RNA sequencing. a) Summary of the direct programming (DP) and standard protocol (SP) differentiation strategies, b-c) tSNE visualization of single cell RNA sequencing data from each differentiation strategy. Timepoints are shown inset. Cell state clusters are color coded and annotated with their identities: ESC = embryonic stem cell; NP = neural progenitor; PNP = posterior neural progenitor; PVNP = posterior and ventral neural progenitor; MNP = motor neuron progenitor; EMN = early motor neuron; LMN = late motor neuron; CN = cortical neuron; DSCN = dorsal spinal cord neuron; Gil = glia type 1; GI2 = glia type 2; Mus = muscle; Meso = mesoderm; Endo = endoderm; Oligo = oligodendrocyte; Astro = astrocyte; Stro = stromal. Arrows inside the bounded area indicate the hypothesized cell state progression during MN differentiation for each method, d-e) Expression heatmap of marker genes used to identify each subpopulation. Colors and annotations for the subpopulations are the same, f-g) Subpopulation abundances for each protocol over time. DP has significantly high MN production efficiency than SP (66% vs. 9%) at the late timepoint. MN = EMN or LMN; NP = NP, PNP, or PVNP; Other = everything else. Colors and labels match b-e).

Our single cell data allowed us to define the efficiency of MN production for each method. For DP, MN production was observed as early as day 4 (19%), and increased over time to 66% by day 11 (Fig. 2F). A minority of off-target neuron subtypes, glia, and miscellaneous cells were also identified. In the SP 23% and 9% of the population resembled MNs at days 5 and 12 respectively (Fig. 2G). This lower efficiency was accompanied by a far larger fraction of off-target products including oligodendrocytes (7.2%), astrocytes (8.6%), muscle (30.6%), and stroma (8.9%) that together accounted for 55.3% of the population by day 12.

### The DP differentiation trajectory lacks intermediates expressing Olig2 and Nkx6-1

What are the differentiation paths taken by each protocol? In the SP differentiation path, cells transit through seven states (Fig. 2C and 2E). These state transitions parallel patterning events in the embryo^13,23,24^: cells first commit to the neural lineage (Sox1+/Sox3+), then are posteriorized (Hoxb8+/Hoxd4+), ventralized (Nkx6-1+/Olig2+), enter the committed MN progenitor state (Mnx1+), and then mature into a neuronal phenotype (Tubb3+/Map2+). This is not a surprise as the growth factor cocktail defining this method was designed to reflect the signaling events taking place in the embryo. By contrast, we found that the path produced by DP was condensed relative to the SP path (Fig. 2B and 2D), consisting of only four states as opposed to seven. After neural commitment (Sox1+), cells immediately began expressing committed MN markers (Mnx1+/Tubb3+), seemingly without the typical spinal embryonic intermediates (Olig2-/Nkx6-1-). A lack of Olig2 expression during DP has been observed previously^3^, and our results confirm at the single cell level that intermediates expressing Olig2 and Nkx6-1 appear entirely absent. Olig2 is necessary for MN development in embryos^1^, indicating the DP drives differentiation from ESC into MNs through a new route.

We confirmed that our inferred dynamics from snapshot single-cell data correspond to the actual underlying differentiation dynamics by performing a dense qPCR time course for a panel of MN genes (Supp. Fig. S3). These bulk measurements confirmed that, for DP, committed MN markers are upregulated immediately following early neural progenitor genes in real time.

### The DP and SP trajectories bifurcate after early neural commitment and converge separately to the MN state

Since the DP omits spinal embryonic intermediates characteristic of the SP path, there must be one of two possible trajectories. Either DP must discontinuously transition from an early neural progenitor into a MN, or it must transit through alternate intermediate state(s). To determine which of these possibilities was the case, we employed a data visualization technique called SPRING^25^ to directly compare the topology of both paths. While tSNE is a powerful method for identifying discrete cell states, SPRING provides a complementary description emphasizing continuum gene expression topologies. SPRING builds a k-nearest-neighbor graph over cells in high-dimensional gene expression space, and then renders an interactive 2D visualization of the cell graph using a force directed layout. This representation revealed that the DP and SP trajectories overlap during early neural commitment, but that they then bifurcate and transit distinct paths that converge independently to the same MN state (Fig. 3A). The dynamics of gene expression over these trajectories resembled the behavior inferred using tSNE, with DP omitting intermediate progenitor genes following its bifurcation from the SP path (Fig. 3B).

**Fig. 3.**
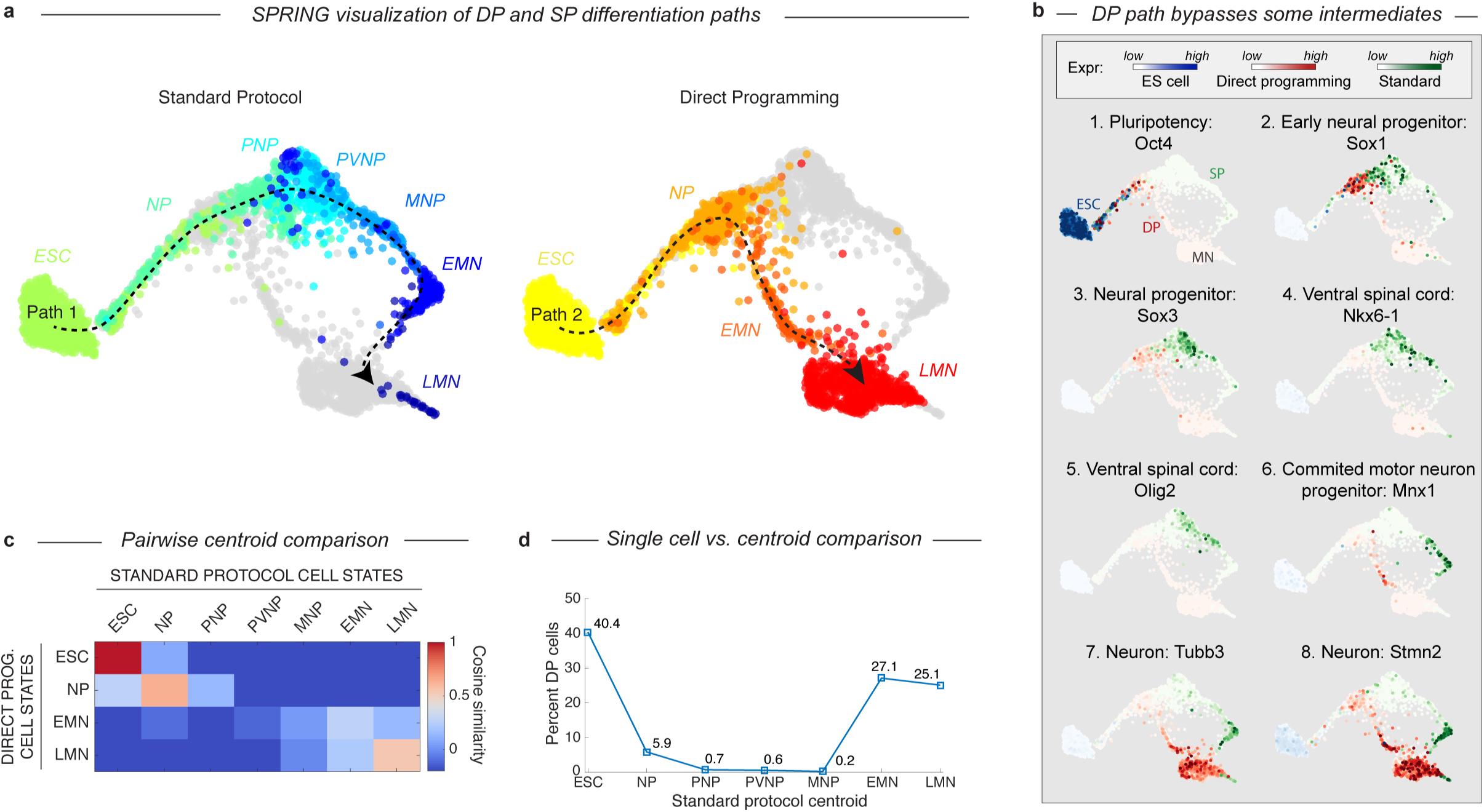
DP and SP induce distinct differentiation paths to the motor neuron state. a) Visualization of differentiation paths for both protocols using SPRING reveals two paths to the same state. DP and SP overlap during early neural commitment, but then bifurcate and converge to the same final MN state separately. Reds = direct programming path; Blues = standard protocol path; arrows indicate hypothezed differentiation trajectories. Cell states are colored and labeled by according to their definition in Figure 1 for comparison, b) Gene expression of key marker genes along each differentiation path confirms exit from the pluripotent state (Oct4) and progression towards the MN state (Tubb3, Stmn2) for both protocols. Only the SP upregulates Olig2 and Nkx6-1, which mark important MN lineage intermediates in the embryo; this occurs following the bifurcation of both paths. Expression in cells from each sample are colored using either a blue (ESC), red (DP), or green (SP) colormap to allow tracking of each path separately, c) Pairwise cosine similarity of cell state centroids. Note early and late similarity of states, but prominent differences during intermediate state transitions, d) Every individual cell of the DP trajectory was assigned to its most similar SP state using a maximum likelihood method. DP cells map to early and late but not intermediate states of the SP cell state progression.

The bifurcation and subsequent convergence of the two differentiation paths can also be appreciated by two other complementary analyses. Pairwise cosine similarities between the cell states from both trajectories (Fig. 3C; Supplementary methods) indicate similarities between the early states (ESC and NP; cosine similarity > 0.64) and late states (LMN; cosine similarity = 0.55), but not the intermediate states (PNP, PVNP, and MNP; cosine similarity < −0.28, −0.09, and 0.06 respectively). We also assigned every individual cell along the DP path to its most similar cluster in the SP path using a maximum likelihood method (Fig. 3D; Supplementary methods). This showed that it was virtually impossible to find a single cell resembling the SP intermediate progenitors in the DP approach. Similarity was again seen only at the early and late states.

### DP transitions through an abnormal intermediate state with forebrain gene expression

The bifurcation of the SP and DP trajectories leads to different intermediate cell states in each case. A total of 26 transcription factors (TFs) are differentially expressed between the DP and SP intermediate states (Fig. 4A). A majority of these (61%) were involved in an anterior-posterior positional gene expression axis. The SP intermediates were enriched more than 6-fold for nine posterior and spinal TFs including Olig2, Nkx6-1, Lhx3, and six Hox genes with a corrected p-value < 0.001. Each of these TFs is expressed in embryonic MNs. By contrast, the DP intermediates were enriched for seven forebrain TFs including Otx2, Otx1, Crx, Six1, Dmrta2, Zic1, and Zic3 at the same stringency, despite the absence of MNs in the forebrain of embryos. Anterior gene expression was previously observed through bulk measurements of DP^3^, and our results reveal that it occurs within a specific subpopulation of cells in the process of differentiating into MNs. We validated these expression differences by isolating intermediate populations of each differentiation path using a Mnx1::GFP reporter cell line, since Mnx1 expression is localized precisely to the distinct intermediate populations of each path (Fig. 3B). Bulk comparisons of these two populations confirmed the enrichment of forebrain TFs, and depletion of spinal progenitor and positional genes in the DP intermediates with just one exception – Zic1 was enriched in DP by our single cell comparison but in SP by microarray (Fig. 4A).

**Fig. 4.**
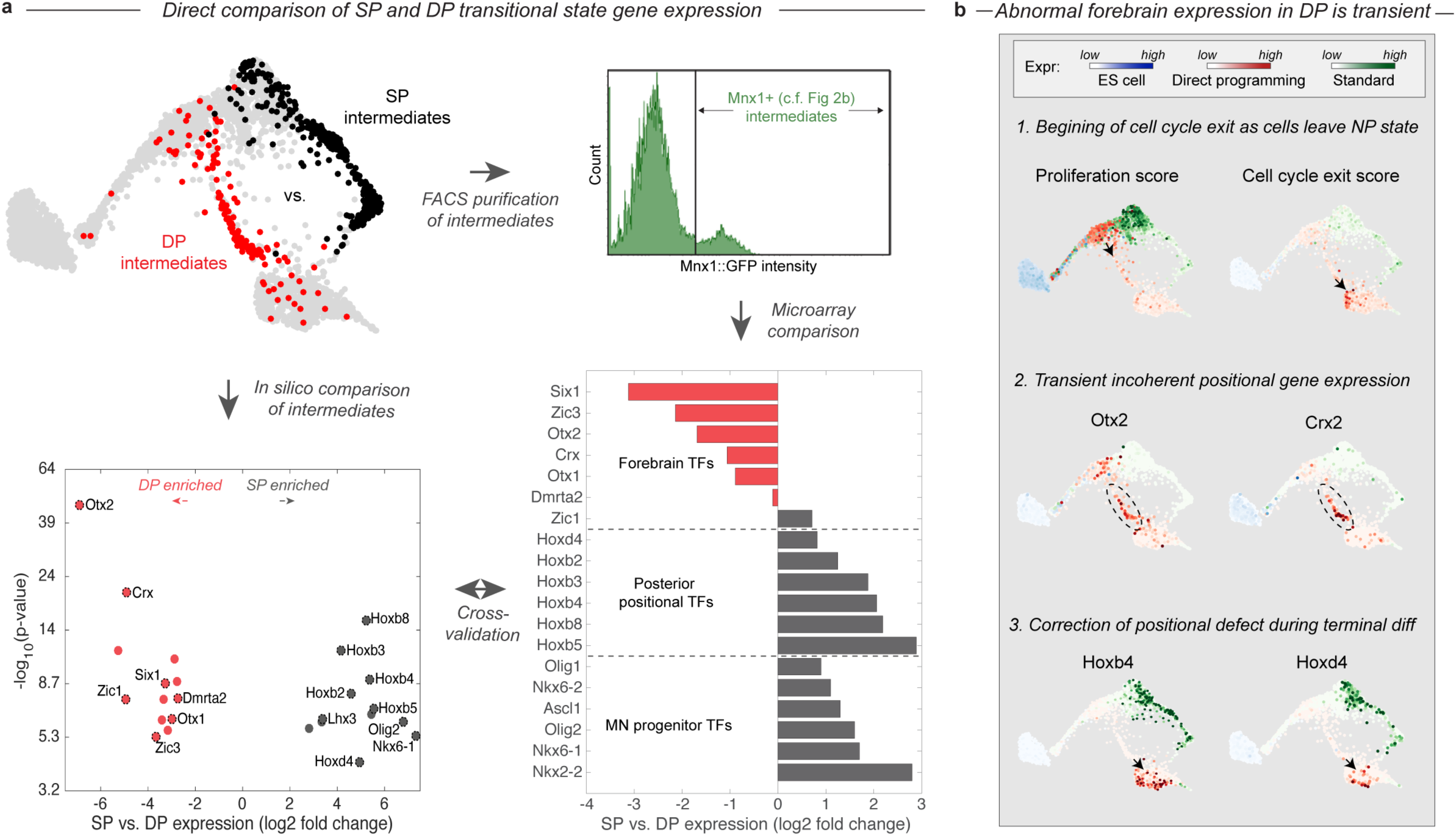
The DP differentiation trajectory involves an abnormal transitional state characterized by a transient positional gene expression defect. a) The DP (red, EMN) and SP (black, PVNP+MNP+EMN) differentiation paths diverge into distinct intermediate states following early neural commitment. These distinct populations were compared by *in silico* differential expression analysis using single cell data (bottom left), or by FACS purification of the intermediate populations using and Mnx1::GFP reporter cell line followed by microarray analysis (bottom right). Fig. 3b. shows that Mnx1 expression localizes to the distinct intermediate populations of both protocols, making it an appropriate target for their purification. Differential expression analysis revealed a total of 26 differentially expressed TFs between the intermediate populations (>6-fold differentially expressed, p<0.001), of which (61%) were involved in either forebrain positional identity (Otx2, Crx, Zic1, Six1, Dmrta2, Otx1, and Zic3) and enriched in DP, or in posterior spinal identity (Hoxb8, Hoxb3, Hoxb2, Hoxb4, Hoxb5, Hoxd4, Lhx3, Olig2, and Nkx6-1) and enriched in SP. The independent microarray comparison confirmed this differences with only one exception - Zic1. It also revealed that the SP intermediates are enriched for additional spinal neural progenitor genes including Ascl1, Nkx6-2, Olig1, and Nkx2-2. b) Before forebrain genes are expressed, DP cells have a high proliferation score and low cell-cycle exit score (computed as the sum of a panel of cell-cycle-associated or tumor supressor genes respectively). They reduce cell cycle gene expression as they enter the abnormal transitional state, and upregulate forebrain genes including Otx2 and Crx. This abnormal forebrain expression is shut off as cells exit the transitional state into the final MN state. The final transition into a MN state is also accompanied by upregulation of posterior positional genes including Hoxb4 and Hoxd3, thus correcting the transient forebrain positional expression defect. Expression in cells from each sample are colored using either a blue (ESC), red (DP), or green (SP) colormap to allow tracking of each path separately.

The abnormal positional gene expression signature that characterizes the DP intermediate state appears transient. Forebrain gene expression is upregulated along the DP differentiation path as cells exit the early NP state into the EMN intermediate state (Fig. 4B). This transition is also accompanied by the downregulation of proliferation-associated genes (Fig. 4B; Supp. Fig. S4). By the time cells exit the EMN state and transition into the more differentiated LMN state, they downregulate forebrain genes and replace this abnormal positional signature with a spinal Hox expression signature characteristic of normal MNs (Fig. 4B). Thus cells converge to the MN state in positional as well as neuronal identity gene expression in the final stages of DP.

### Both DP and SP approach a transcriptional state similar to *bona-fide* MNs in embryos

Given that the two protocols induce distinct – and in the case of DP, unnatural – differentiation paths, we were curious how their final products compared with primary MNs. We harvested MNs from the embryo of a Mnx1::GFP reporter mouse and performed inDrops measurements on 874 E13.5 MNs after FACS purification. Though the majority of Hb9+ sorted cells were MNs (73.8%), this population also contained glia (20.1%), fibroblast-like cells (1.8%), and immune-type cells (1.2%; Fig. 5A; Supp. Fig. S5). Using only the cells identified as *bona-fide* MNs, we probed the similarity of the *in vitro* derived cells to the primary MNs, using three measures: global similarity of the transcriptomes (cosine similarity); co-clustering frequency; and differential gene expression analysis. In both paths, neurons become more similar to primary MNs over time (Fig. 5B). The clusters most highly correlated with primary MNs were the LMN state from the DP protocol (cosine similarity = 0.62), and the LMN state from the SP (cosine similarity = 0.37). Notably, only 2.7% of output cells from the SP were in the LMN state, compared to 62.7% for DP. Thus, the ratio of the efficiency of forming LMNs by the DP protocol to the SP protocol is 23 fold, seven-fold higher than what we calculated based on a comparison of marker genes alone. At the level of single cells, co-clustering of the different experiments showed that 95% of DP MNs robustly co-cluster with primary MNs compared with 26% for SP derived MNs (Supp. Fig. S6). However, differential gene expression analysis revealed that neither protocol perfectly recapitulates the gene expression profile of MNs isolated from the mouse. Both protocols showed a depletion of the most posterior Hox genes, perhaps indicating an anterior spinal cord identity of *in vitro* MNs, and a small enrichment of several genes related to microtubule function and cell cycle exit that may indicate subtle differences in neuronal maturation (Supp. Fig. S7).

**Fig. 5.**
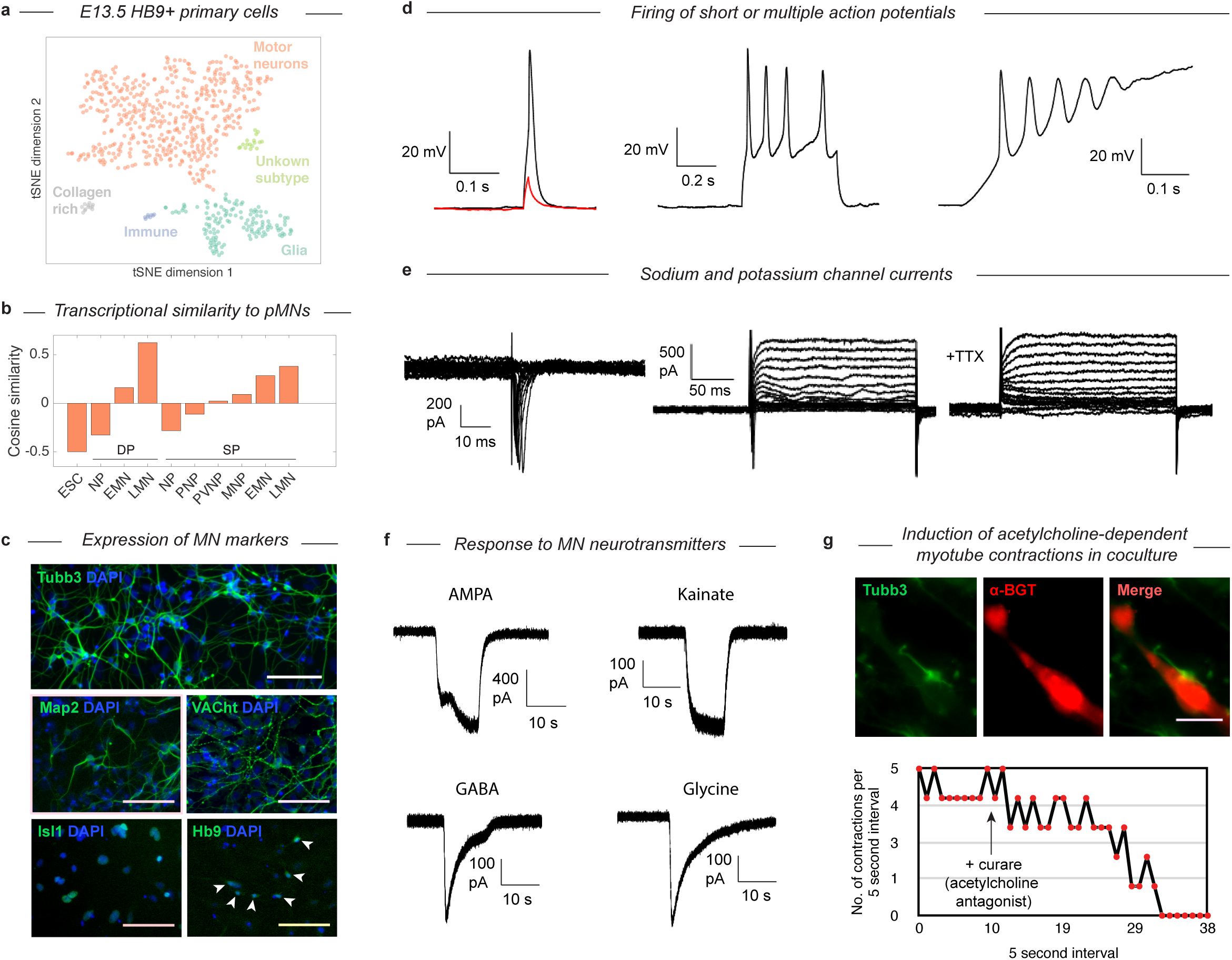
Validation that DP and SP trajectories both approach a *bona-fide* MN state. a) tSNE visualization of 874 single cell transciptomes from FACS purified Mnx1+ MNs from embryos reveals heterogeneity within this population. b) Comparison of cell states along both trajectories with true primary MNs (orange population in a). In both DP and SP similarity increases as differentiation proceeds. Late DP states are the most similar to embryonic MNs. c) Immunostaining of MN markers in DP MNs confirming expression and correct subcellular localization of Tubb3, Map2, VACht, Isl1, and Hb9. DP MNs also: d) can fire single or multiple action potentials upon stimulation, e) show sodium and potassium channel currents, and f) are responsive to multiple MN neurotransmitters - AMPA, kainate, GABA, and glycine. g) Co-culture experiments show that DP MNs can induce clustering of acetylcholine receptors on muscle myotubes (indicated by α-BGT staining) and induce their contractions. The induced contractions are dependent on MN activity as they can be blocked by addition of the acetylcholine antagonist curare.

### DP MNs have structural and functional properties of true MNs

Having established that MNs derived via both gene expression trajectories reach roughly the same MN transcriptional state, we wished to validate that their function and structural organization was also independent of their distinct developmental histories. The SP has been characterized extensively as giving rise to functional MNs^13^, so here we examined structural and functional characteristics following DP. We confirmed that selected protein content matches the mRNA markers by immunostaining for Tubb3, Map2, VACht, Isl1, and Hb9 (Fig. 5C). Tubb3 and Map2 were present, and VACht was seen at discrete puncta on the axons (suggesting localization to acetylcholine secretory vesicles). TFs Isl1 and Hb9 were localized in the nucleus. Finally, the GFP from the Mnx1::GFP reporter was activated and expressed in the cytoplasm. To test the functional properties of the DP MNs, we performed whole-cell patch clamp recordings. Depolarization induced single or multiple action potentials in current-clamp experiments (Fig. 5D). Depolarizing voltage steps induced fast inward currents and slow outward currents characteristic of sodium and potassium channels, respectively (Fig. 5E). Exposure to 500 nM Tetrodotoxin (TTX) blocked the inward current, indicating sodium channel involvement. We then tested whether our DP neurons would respond to neurotransmitters that act on MN (Fig. 4F). Exposure of the neurons to AMPA, kainate, GABA, and glycine (100 μM each) induced in each case inward currents similar to that seen in primary embryonic MNs. To see if the DP neurons could also form neuromuscular junctions, we co-cultured the neurons with differentiated C2C12 skeletal muscle myotubes and incubated them for 7 days. We observed clustering of acetylcholine receptors on the C2C12 myotubes near contact points with the DP neurons, which can be seen with alpha-bungarotoxin (α-BGT), which binds to acetylcholine receptors (Fig. 4G). We then observed regular contractions of some C2C12 myotubes that began after several days in co-culture (Fig. 4G, Supp. Video 1). These contractions could be stopped by the addition of 300 μM Tubocurarine (curare), an antagonist of acetylcholine receptors, indicating that the contractions were induced by the acetylcholine release from the MNs. Similarly, we noticed that the DP MNs could induce contractions in DP muscle myotubes that we previously generated with MyoD (Supp. Video 2)^4^. These results confirm that DP MNs have the expected functional properties of *bona-fide* MNs.

## Discussion

The results we have described provide evidence at the single cell level that differentiation can proceed by multiple routes yet converge onto similar transcriptional states. We show that while using the SP cells are driven to retrace the embryonic lineage, DP induces cells to differentiate through a dramatically different path. The DP path bypasses multiple intermediate progenitor-states that evolved in the embryo, and yet still converges to the same discrete and recognizable MN phenotype. This convergence occurs from an abnormal intermediate state, and does not appear to involve a shared set of terminal cell state transitions; it is highly orthogonal. Moreover, as cells converge they manage to not only establish gene expression related to MN functions, but they also correct positional gene expression defects (exchanging forebrain for spinal gene expression) in the absence of external signals. Relative to our initial research questions (Fig. 1), we conclude that DP of mESCs into MNs occurs via a late bypass that involves alternative intermediate states not seen in the embryo, and that this new route converges near perfectly to the same final state. Convergence into a MN therefore does not appear to depend rigidly on the precise history of intermediate states through which cells differentiate.

This ‘history independence’ of the final state is consistent with a dynamical view of gene regulation in which cell states correspond to ‘attractor basins’, i.e. stable states of gene expression that are robust to modest perturbations. If attractor basins do not exist, the precision of the observed overlap between DP and SP MNs would require a special coincidence, like finding a needle in a high dimensional haystack. The concept of cell states behaving as attractors has been proposed previously to explain several properties of blood cell types^26-28^. There are at least two important corollaries of this behavior applying in development. From a practical perspective, it is a common concern that DP methods may generate cell types with subtle defects due to their unusual developmental histories^9^. Attractors would be robust to this vulnerability and indeed our results show that it is not necessary to recreate the precise sequences of steps taken in embryos to generate *bona-fide* MNs. It could also hint at a mechanism that might help animal body plans evolve flexibly. Specifically, by decoupling the identity of mature cell state attractors from their developmental histories evolution would be able to act on each independently. In principle this could contribute to evolvability by allowing mature cell states to be transposed onto new lineages in new body locations.

The mechanisms that define the MN attractor basin and allow the artificial DP trajectory to converge onto the correct final state are largely unclear. The MN state is thought to be stabilized by a network of self-reinforcing TFs^24^, involving Mnr2, Mnx1, Lim3, Isl1, Isl2, and Lhx3, Ngn2, Myt1l, Nefl, and Nefm. DP aims to kick-start this network by activating a subset of important components. Yet, far from immediately activating this network, our data show that DP initially drives cells to differentiate into an early NP state through the same pathway as the SP trajectory, seemingly oblivious to the DP TFs, and then even activates non-MN genes in the transitional state. Understanding why the activation of the MN program lags behind TF induction may provide important clues into how the DP factors act. One possible source for a lag is that activating a complete neuronal program requires first activating additional core TFs (so-called ‘feed-forward’ circuitry). Indeed, recent studies have shown that Ebf and Onecut are activated by Ngn2 (one of the DP TFs), and that both are required to subsequently direct binding of Isl1 and Lhx3 (the other two DP TFs) to MN target genes across the genome during DP^12,29^. A second possible source of lag is that extracellular signaling provides inputs that immediately affect cell state, but take time to sensitize cells to the DP factors. For example, signaling changes might activate DP TFs through post-translational modifications, by activating co-factors, or by inducing chromatin state changes. We have indeed observed that MNs are not generated if DP TFs are induced in cells cultured in pluripotency media, indicating a requirement for changes in signaling (not shown). Conversely, when mES cells are transferred to minimal media without inducing DP factors, they acquire a forebrain neural progenitor identity by default^30-32^. This suggests that the early dynamics and abnormal forebrain / MN expression of the DP transitional state might in fact be driven by the signaling environment and not the DP TFs. These alternatives suggest future experiments to better resolve the mechanisms driving the DP, by re-mapping the trajectory induced during DP after changing signaling conditions, or the choice of DP TFs.

As a methodology, DP is significantly more efficient than the SP without loss of quality in the MN populations produced (Fig. 2F-G; Fig. 5). The high-efficiency of DP likely derives from both its more uniform experimental conditions as well as its more direct differentiation path. Experimentally, DP: relies on 2D rather than 3D tissue culture (as in the SP), minimizing uncontrolled cell-cell communication; forces every individual cell to express MN TFs from a genetically integrated construct, increasing uniformity; and employs a defined-media without growth factors that may minimize proliferation of progenitor states. The more direct differentiation path induced by DP should also itself increase MN conversion efficiency by minimizing error propagation through chained opportunities for off-target fate choices. During sequences of intermediate cell state transitions, each transition can have competing off-target fates. Thus, differentiation processes that involve many sequential intermediate transitions suffer from multiplicative efficiency losses. Indeed, the longer sequence of intermediate states the in SP generates a far larger fraction of off-target populations that increases with time, suggesting a progressive loss of efficiency. Targeting terminal attractor basins through the shortest possible differentiation paths may prove to be a generally effective strategy to generate desired cell states.

## Acknowledgements

We are grateful to Esteban Mazzoni for providing the transcription factor cassette containing ES cell line used in the DP experiments presented here.

## Competing interests

V.L. and M.K. are cofounders of StemCellerant, LLC. A.K. and M.K. are cofounders of 1CellBio, Inc.

## Author contributions

All authors helped design the study and its experimental questions. V. L. established the MN differentiation protocols and performed their molecular characterization. J. B. performed the single-cell data collection and analysis. S.L. did the electrophysiology recordings. M. K., A. K. and C.W. assisted with analysis and interpretation of experimental results. All authors contributed to the writing of the manuscript and preparation of figures.

---

**Fig. S1.**
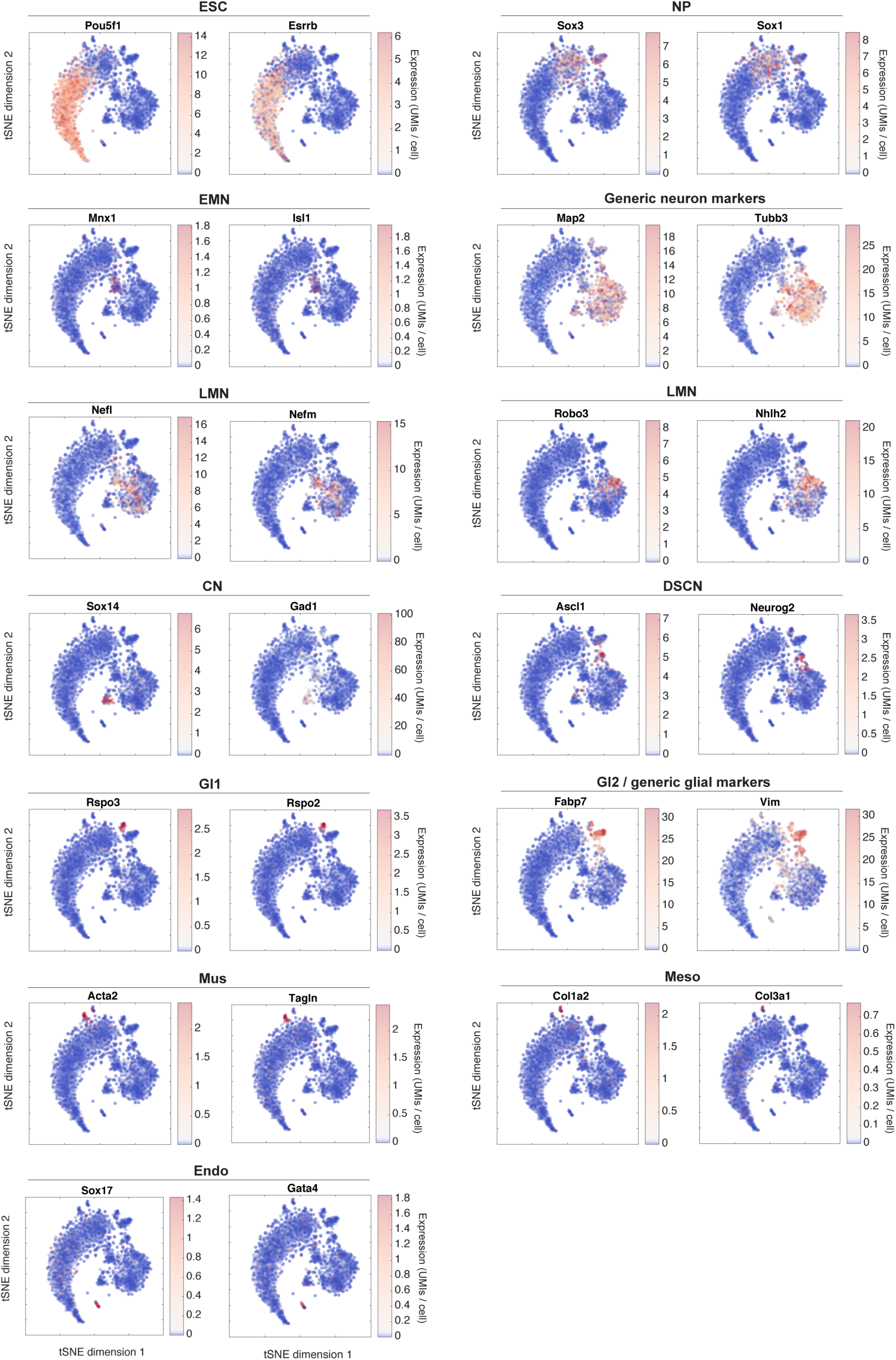
tSNE visualization of marker genes for DP subpopulations. This figure relates to main text Fig. 2. It shows the raw expression of sample marker genes that were used to identify each subpopulation. Two marker genes are shown per state.

**Fig. S2.**
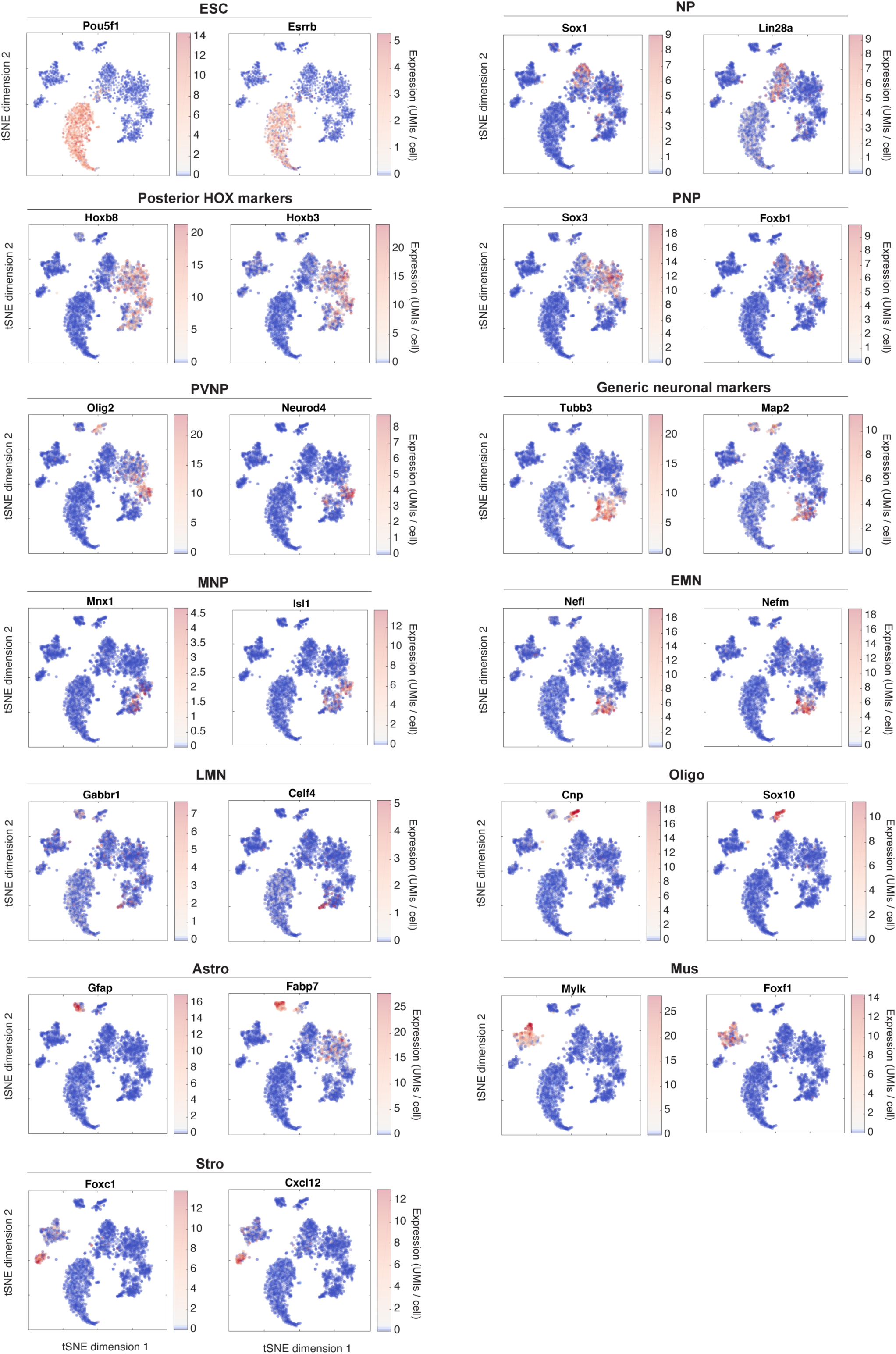
tSNE visualization of marker genes for SP subpopulations. This figure relates to main text Fig. 2. It shows the raw expression of sample marker genes that were used to identify each subpopulation. Two marker genes are shown per state.

**Fig. S3.**
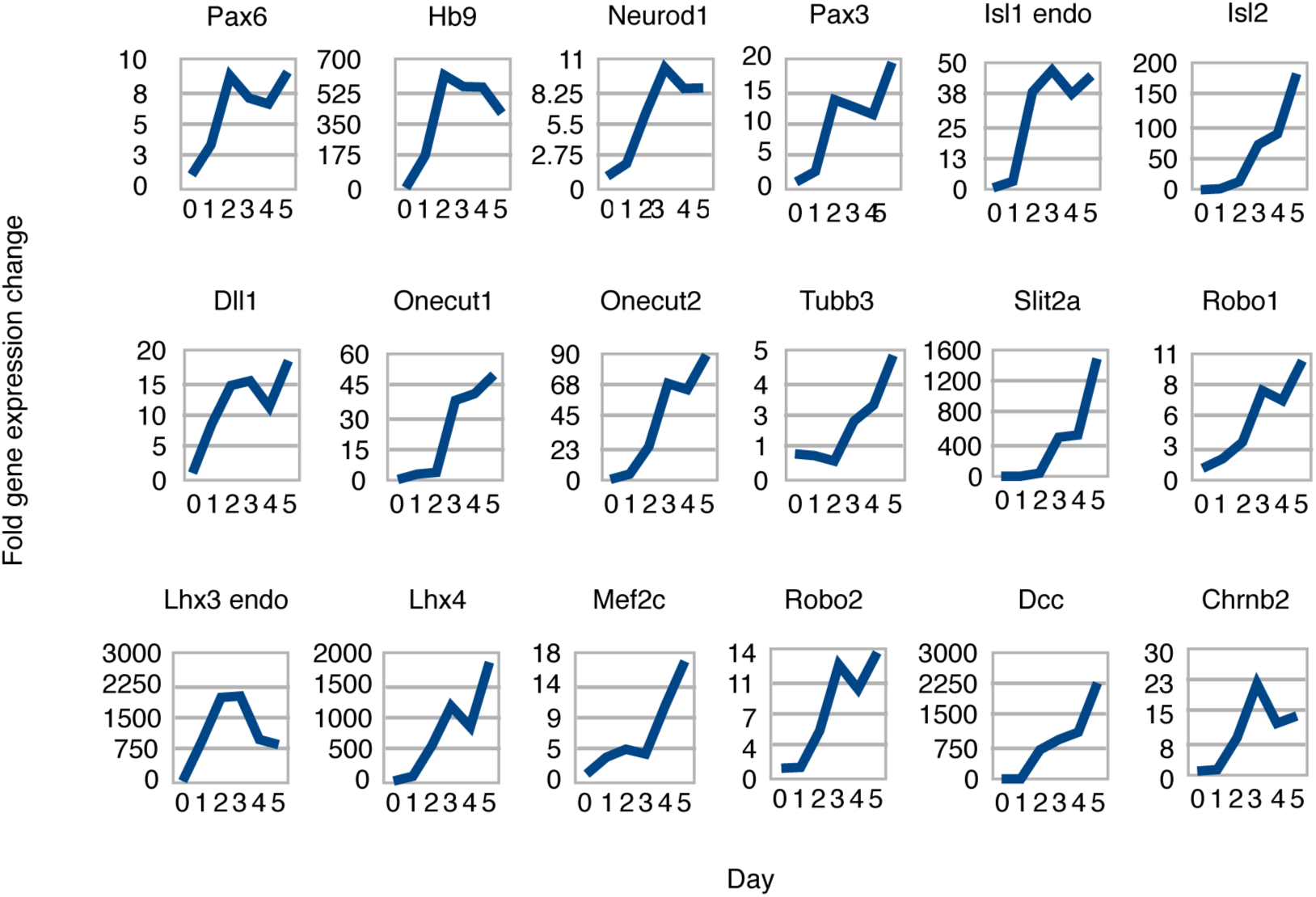
qPCR validation of inferred MN differentiation gene expression dynamics. Measurement of the bulk population gene expression over time helps to confirm that the trajectory inferred from single-cell analysis matches the true dynamical events of our system. These results confirm that key MN differentiation markers are induced on a real timescale that matches their ordering in our Inferred trajectory. Terminal genes such as Mnx1 and Tubb3 are upregulated Immediately after Induction of early progenitor genes such as Sox1. The values plotted are shown In units of fold change relative to their expression In day 0 mESC cultures.

**Fig. S4.**
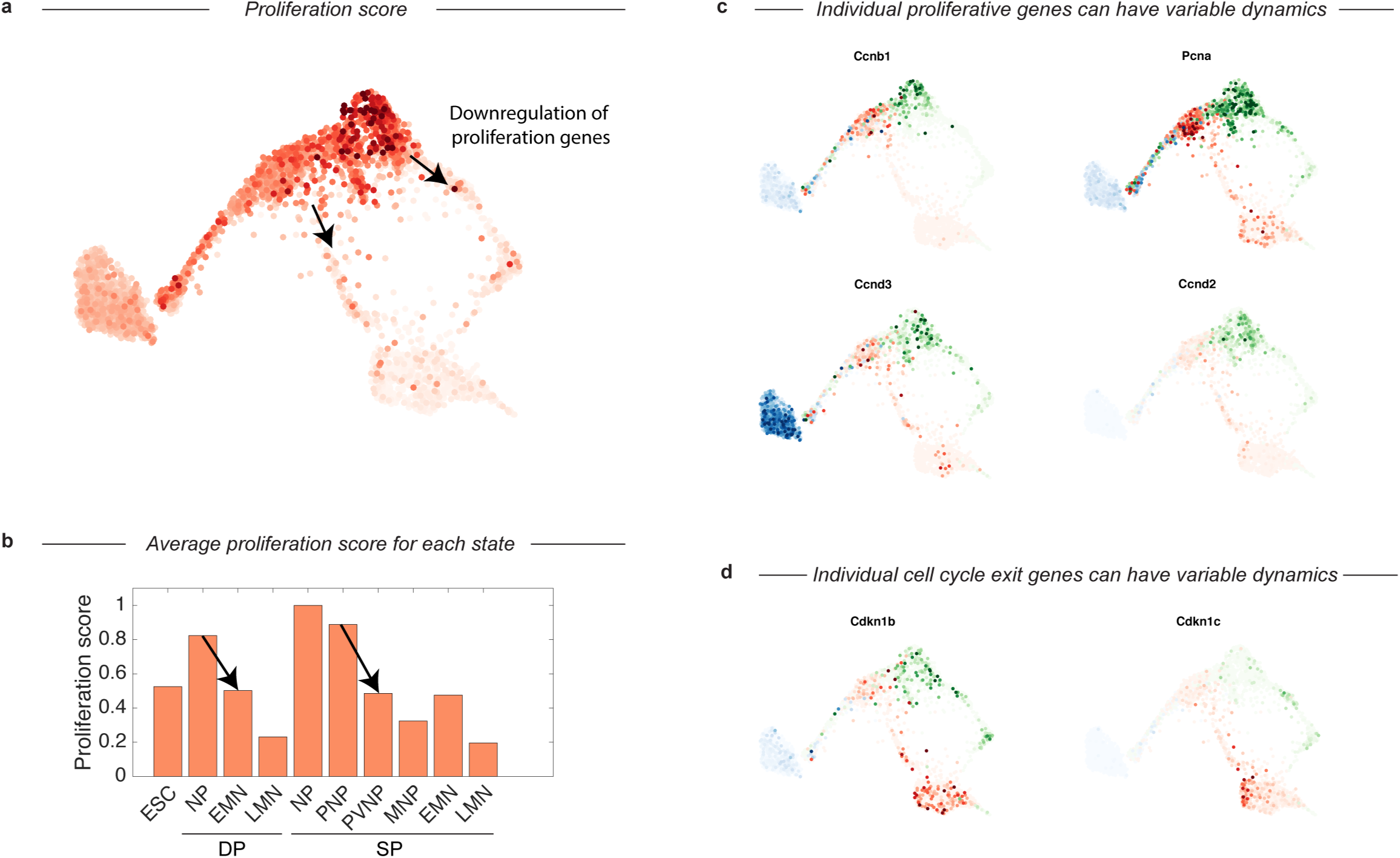
Cell-cycle gene expression is restricted to early progenitor states. This figure relates to main text Fig. 4B. a) Cell cycle gene expression score overlayed on SPRING trajectory shown in main text. The score is the summed expression of a panel of 21 cell-cycle related genes. Cell cycle expression is high in ES cells and neural progenitors from both protocols, but drops sharply upon entry into the transitional states preceding terminal differentiation.b) Shows the average proliferation score along both DP and SP trajectories computed by state (and max normalized). Arrows indicate the same transition as the arrows shown above in panel a. c-d) Example raw expression data for representative individual cell cycle or tumor suppressor / cell-cycle exit genes. Some genes appear to be enriched in specific states, such as Ccnd3 in ES cells and Ccnd2 in neural progenitors from the SP. Expression in cells from each sample are colored using either a blue (ESC), red (DP), or green (SP) colormap.

**Fig. S5.**
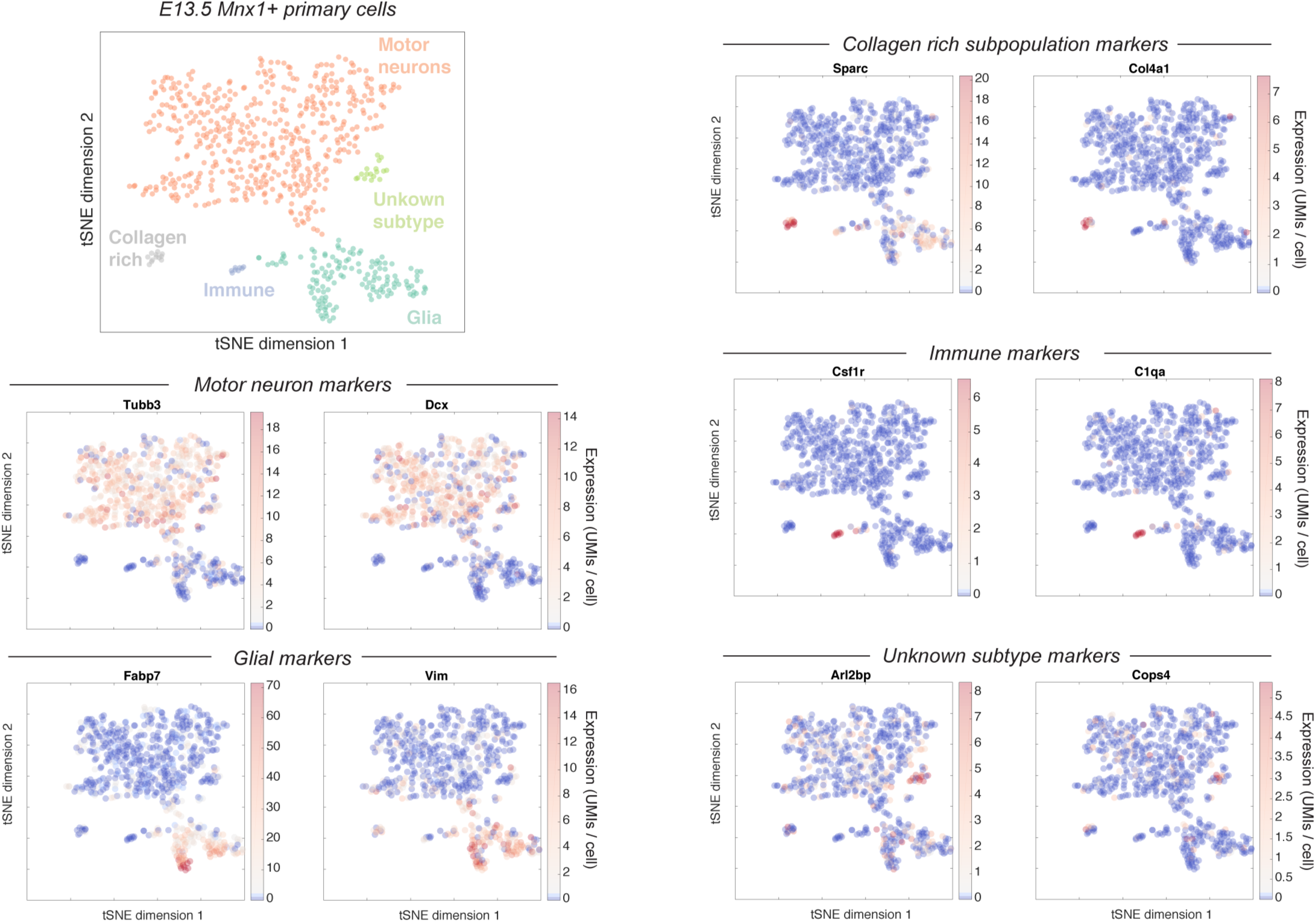
Heterogeneity within the Mnx1+ E13.5 primary motor neuron population. This figure shows the raw expression of a set of marker genes that were used to identify the subpopulations indicated in main text Fig. 5A. Two markers are shown per subpopulation, with their expression intensity overlaid on the original tSNE embedding.

**Fig. S6.**
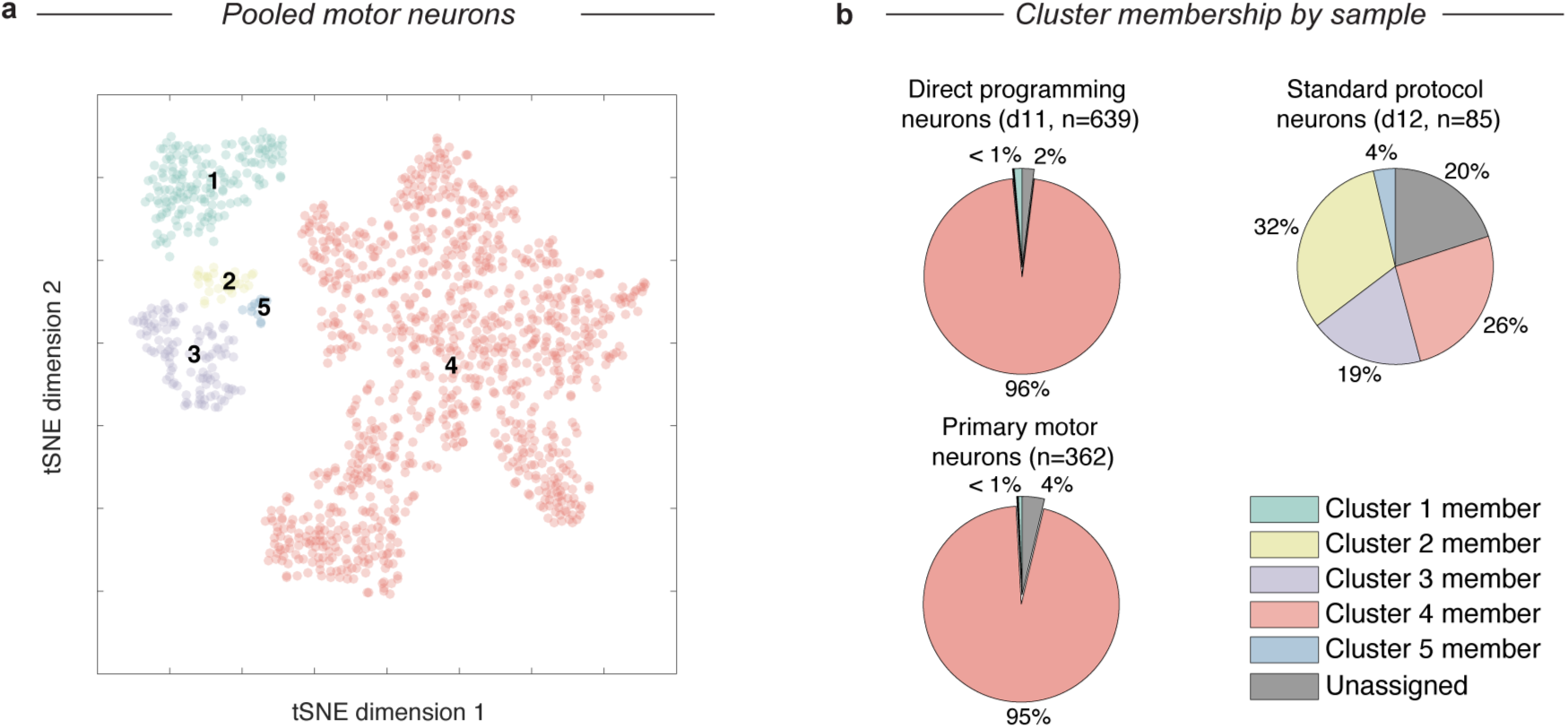
Co-clustering analysis of MNs from embryos versus DP and SP MNs. a) Neuron populations (LMN) from both protocols were pooled with Mnx1+ FACS purified E13.5+ motor neurons (pMNs), visualized using tSNE, and clustered using density-gradient clustering, b) For each of DP, SP, and pMNs, the percentage of cells belonging to each cluster in a) was calculated. 95% of both DP MNs and pMNs group together in cluster 4, indicating a high degree of similarity, while MNs from the SP are more mixed in their memberships. Pie chart colors and cluster numbers match a). Note that unassigned cells are not plotted in a).

**Fig. S7.**
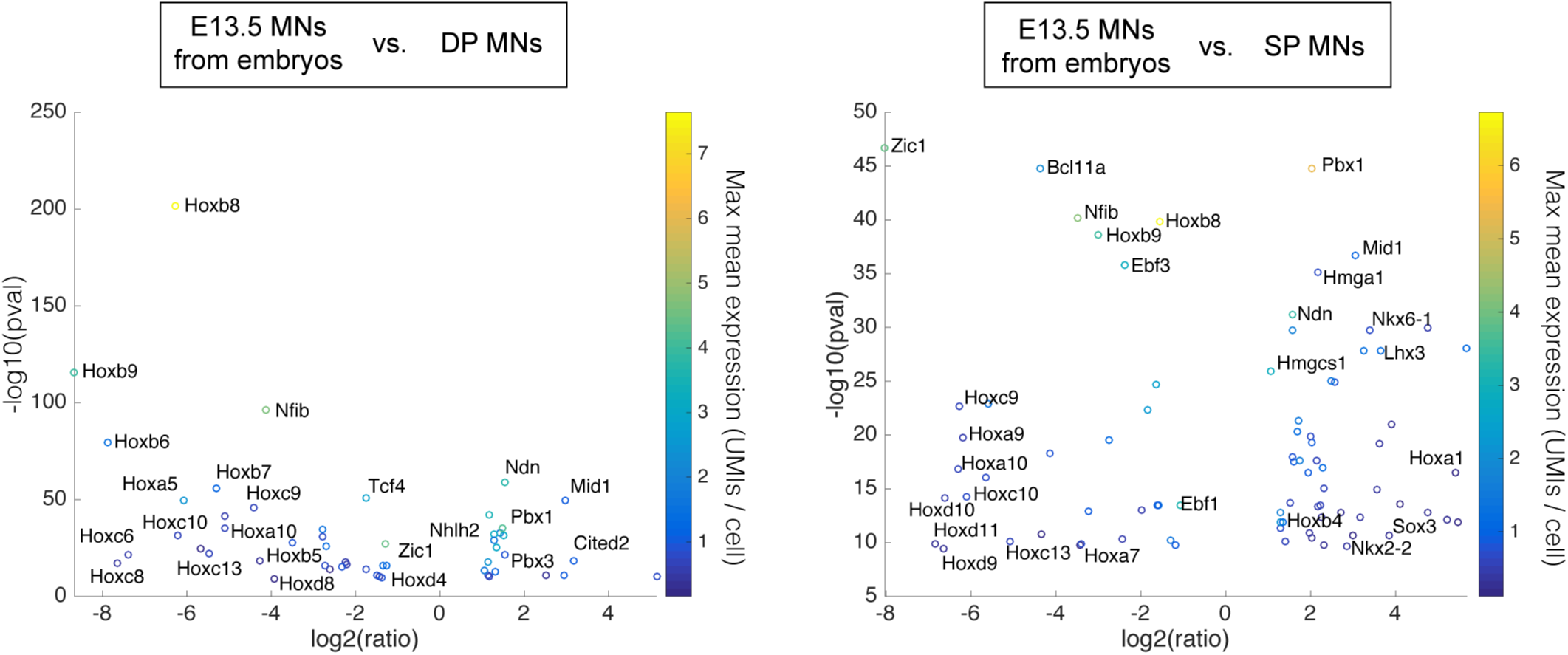
Differential gene expression between DP or SP MNs and E13.5 MNs from embryos. This figure relates to Fig. 5. of the main text. It shows the transcription factors that are differentially expressed between the final MN states of each protocol and E13.5 Mnx1+ FACS-purified MNs from the embryo, computed from single cell expression data. The dominant transcriptional signature for both protocols is a depletion of the most posterior Hox genes (Hox*7 and higher), indicating that both DP and SP MNs may have a trunk or more anterior (not hindlimb / tail) identity. Overexpressed genes common to both protocols include a small number related to neuronal maturation such as Mid1, Ndn, Pbx1, and Pbx3. Each panel is a volcano plot for the indicated comparisons. Only genes with a corrected p-value < 001 and an expression ratio >2 are shown. A colorbar is used to indicate the highest mean expression of each gene in the states being compared.

## Supplementary Methods

### ESC culture

The mouse ES cell line containing doxycycline-inducible Ngn2+Isl1+Lhx3 (NIL) and the Hb9::GFP reporter was graciously provided by Esteban Mazzoni. ESCs were cultured in standard media (DMEM with LIF + 15% fetal bovine serum) on 0.2% gelatin-coated dishes.

### Differentiation into motor neurons: direct and standard programming

Twenty-four hours before starting differentiation, ESCs were trypsinized and seeded onto plates precoated with a mix of poly-d-lysine (100 μg/ml) and laminin (50 μg/ml) instead of gelatin for adherence. ESCs were counted by a Beckman Coulter Counter and seeded at a density of approximately 200,000 cells per well of a 6-well dish. At day 0 (ESCs), the media was switched from standard ES media to N2B27 media (Invitrogen). Doxycycline was also added at 3 μg/ml starting to induce expression of the NIL transcription factors. Media was changed daily. For the standard programming protocol, the steps described in Wu et al. were followed strictly.

### Mouse embryonic motor neuron cultures

The B6.Cg-Tg(Hlxb9-GFP)1Tmj/J mice (JAX# 005029) were bred with C57BL/6J (JAX# 000664) for embryonic motor neurons dissection. All animal protocols were approved the Institutional Animal Care and Use Committee at Boston Children’s Hospital. On gestational day 13 (E13), the female mice were anesthetized and all embryos were collected by caesarian section. Only GFP embryos were used for further dissection. The spinal cords were isolated and their meninges were removed. Each isolated spinal cord was dissociated by trypsin and mechanical trituration. After filtering the cells with 100 μm strainers, the cells were spun down and re-suspended in PBS, and subjected to flow cytometry. Cells were run through a 100 μm nozzle at low pressure (20 p.s.i.) on a BD FACSaria II machine (Becton Dickinson, USA). A neural density filter (2.0 setting) was used to allow visualization of large cells.

### Single cell transcriptomics using InDrops

We dissociated differentiating mESC cultures using a 0.25% Trypsin 2mM EDTA solution (Gibco). Primary HB9+ sorted motor neurons were dissociated as above. Dissociated cell suspensions were verified to be monodisperse and of viability >95% using a coulter counter with trypan blue staining (BioRad). We then performed droplet-based barcoding reverse transcription (RT) reactions and prepared massively multiplexed sequencing libraries using InDrops as described in Klein et al. Briefly: cells, lysis and RT reagents, and barcoding primers attached to hydrogel beads are combined in nanoliter sized droplets suspended in an oil emulsion using the InDrops microfluidic device. A barcoding RT reaction is then carried out in the droplet emulsion, uniquely labeling the RNA contents of every cell using a cell barcode and unique molecular identifier (UMI). Following RT, barcoded emulsions are split into batches, and the emulsion is broken. Combined material is amplified into a nanomolar Illumina sequencing library through a series of bulk reactions: second strand synthesis, in vitro transcription (IVT), fragmentation, RT2, and a final low cycle number PCR. The majority of the amplification takes place during IVT ensuring uniform library coverage. Single cell libraries were then sequenced on either the Illumina HiSeq or NextSeq platforms. Reads were demultiplexed using an updated version of the custom bioinformatics pipeline described in Klein et al˙. A python implementation of this pipeline is now publically available on GitHub. Briefly, it filters for reads with the expected barcode structure, splits reads according to their cell barcode, aligns them to a reference transcriptome (we used GRCm38 with some added mitochondrial genome transcripts), and then counts the number of different UMIs appearing for each gene in each cell. The final output is a counts matrix of cells vs. genes that we loaded into MATLAB for further analysis.

### Single-cell data clean up: minimum expression threshold, total count normalization, stressed cell removal

Before performing the analyses described below, three steps were taken to ensure that the data was of high quality. First, we required all cells to have at least 1000 UMIs detected. This removed any signal potentially coming from empty droplets. Second, data were total count normalized to ensure differences between cells were not due to technical variation in mRNA capture efficiency or cell size. Finally, cells that had a high stress gene signature were excluded from analysis. Stressed cells were initially recognized as a small percentage of cells (<10%) that clustered apart from everything else and specifically expressed very high levels of a mitochondrial gene set that is associated with cellular stress. Masks that convert raw counts into our filtered set are provided online.

### Visualization of single-cell data using tSNE

To visualize high-dimensional single cell data, dimensionality reduction is essential. We chose to implement tSNE as described in Klein et al. The core steps are summarized as follows. Steps 1-2 preceed tSNE, and focus the algorithm on genes that best describe differences between cell populations.

1. Extract the top 1000 highly variable genes. We do this using a statistical test derived specifically for InDrops data (Klein et al.).
2. Extract principal variable genes. Principal variable genes are a subset of the highly variable genes from step 2 that we find best describes the cell population structure. The steps to find principal variable genes are:

a. Perform PCA using the top 1000 biologically variable genes.
b. Identify the number of non-trivial principal components. This is done by comparing the eigenvalue of each principal component from a. to the eigenvalue distribution for the same data after being randomized. Only principal components with eigenvalues higher than those observed on random data are retained.
c. Extract genes that contribute most highly to these principal components by imposing a threshold on the gene loadings for each non-trivial PC.
3. Perform tSNE. We used the MATLAB implementation of tSNE from Van de Marteen et al. As input, we supplied z-score normalized principal variable genes, and asked tSNE to perform an initial PCA to a number of dimensions equal to the number of non-trivial principal components in step 4. tSNE then takes cells embedded in this PCA space and nonlinearly projects them onto two dimensions for visualization.

### Subpopulation analysis

The first goal of our single cell analysis was to describe the identity and proportions of cell states generated by each of two motor neuron differentiation strategies. For this purpose we found that good results were achieved using local-density gradient clustering on the 2D tSNE representation of the data. This approach provided a clear and natural cell state classification that was well aligned with prior knowledge about marker gene expression domains. The steps we took are summarized as follows:

1. Perform tSNE (as above) on cells pooled from all timepoints for each protocol (e.g. Fig. 2B).
2. Apply local density gradient clustering to define cell states.
3. Identify genes specifically expressed by each cell state, and use prior knowledge on their expression domains to generate a cell type annotation (e.g. Fig. 2D).
4. Quantify the fraction of cells in each state at each timepoint (e.g. Fig. 2F).

We identified genes that were specifically expressed by each subpopulation through pairwise t-tests and visually inspected their expression over the tSNE embedding. In Figure 2D-E of the main text, we show the z-score normalized expression of a selection of marker genes that we used as a heatmap. Z-score normalization preserves differences between populations while putting the expression level of every gene on the same scale. In Supplementary Figs. S1 and S2, we show the un-normalized expression of each of these genes individually so that the reader may compare.

### Initial identification of differentiation trajectories

During our subpopulation analysis we observed, for both protocols, a continuous progression of cells that spanned the initial ES cell state through the early and late differentiation timepoints. This progression was punctuated by familiar progenitor states that were ordered in a way that was consistent with known events in motor neuron differentiation (Figure 2D-E), and was correlated with chronology (Figure 2B-C inset). We interpreted these progressions as differentiation trajectories. Each is reconstructed from three population snapshots (day 0, day 4/5, and day 11/12). Because differentiation *in vitro* is asynchronous, the single cell data overlapped from one timepoint to the next. We deduce from this that intermediate states were not missed due to the spacing of our timepoints. We also validated that important intermediate genes were not detected over a densely sampled timecourse using qPCR (see below).

### Differential gene expression analysis between cell states from single-cell data

We identified differentially expressed genes between cell states by using two-tailed t-tests with a multiple hypothesis testing correction. We defined differentially expressed genes conservatively at a FDR of 5% and a significance level of p<0.0001. We only considered genes where at least one of the states being compared had >=10 cells with non-zero expression. To find marker genes of a population we asked for genes that were enriched in that population versus everything else. For several comparisons we restricted our analysis to Riken transcription factors. This list contains ^∼^1,500 genes with manually annotated transcription factor activity. We represented differential expression data using volcano plots, and colored the expression intensity of each gene using a colorbar; the mean of the higher expressing state was used.

### Comparison of differentiation paths I: Visualization of alternate differentiation trajectories using SPRING

After our initial identification of the differentiation paths for the standard protocol and for direct programming we wished compare their routes. One of the most powerful ways to begin addressing such a problem is to simply visualize the data. Yet, we found tSNE gave unclear results for a direct comparison of the paths; visualizing both protocols together resulted in some mixed clusters, and some distinct clusters, but no overall coherence to the representation as we had found looking at one or the other trajectory separately. The limitations of tSNE for analyzing continuous processes are well known.

We therefore turned to a new method developed in parallel to this work in our lab called SPRING, that in our experience does better in analyzing continuous processes in single cell data. SPRING has three core steps: first, it reduces dimensionality to a 50 dim PCA space; second, it constructs a k-nearest-neighbor (kNN) graph in this space; finally, it renders an interactive 2D visualization of this kNN graph using a force directed layout. In this visualization edges of the kNN graph are literally springs that pull together similar cells, while every cell has a magnetic repulsion force that pushes it away from other cells and an intrinsic gravity. The balance of these forces illuminates the topology of how cells are positioned in high-dimension with respect to one another. In this visualization nodes can be moved around to rotate the projection and find the clearest representation; in each new position the graph re-relaxes according to the underlying physics of the force directed layout. The specific steps we took to generate Fig. 3A are as follows. Note that a small manual correction was made to the position of the ESC and LMN populations to reduce white space, but did not distort relationships between populations.

1. Load the cells along each differentiation path into SPRING separately.
2. Remove doublets. Doublets were identified by three criteria. 1) they are rare, in line with our experimental expectation for cell doublets, 2) they formed long range connections between large cell groups in the same timepoint and sample, 3) they do not possess any unique marker genes; all genes they express are a linear combination of two other cell states. We identified approximately 80 doublets in the standard protocol (<3% of all cells), and approximately 40 in the direct programming approach (<2% of all cells).
3. Load the filtered cells from both protocols into SPRING together. To make plots the coordinates from the SPRING representation were exported into MATLAB; cells were either colored by cell state (Fig. 3A), or gene expression (Fig. 3B).

### Comparison of differentiation paths II: Pairwise cosine similarity of cell state centroids

We also asked how the direct programming and standard motor neuron differentiation paths were related to each other by performing a pairwise comparison of cell state centroids. Centroids are the average or center of mass in gene expression space for a collection of cells. Centroids can provide a very accurate estimate of global gene expression that averages over the noise intrinsic to single-cell RNA sequencing at the level of individual cells. By computing the cosine similarity between two cell state centroids we are asking how similar these states are in average gene expression. If the paths do indeed split and then reconverge our expectation was to see similarity between early and late states in both progressions, but not between the intermediate states. We chose to perform these comparisons in a PV-gene space constructed from cells in both protocols to make our comparisons as sensitive as possible. We obtained similar but less sensitive results using all genes above a minimum expression value (not shown). A summary of the steps of this analysis is:

1. Combine all cells from both protocols, extract PV-genes (as described above), and z-score normalize their expression.
2. Calculate the centroid of every cluster from each trajectory in this PV-gene space.
3. Compute the pairwise cosine similarity for all direct programming clusters versus all standard protocol clusters.
4. Visualize: we chose to use a heatmap (Fig. 3C).

### Comparison of differentiation paths III: Maximum likelihood assignment of single cells to cell state centroids between trajectories

Finally, we performed an independent comparison of the differentiation paths that did not depend on our definition of cluster boundaries in the direct programming trajectory. We asked whether any potentially rare individual cells or subpopulations in the direct programming trajectory resemble the intermediate states of the standard trajectory. Because we were now dealing with single cells, not averages of many measurements, care was necessary in this analysis to remain robust relative to the noise in single cell measurements. We therefore took a Bayesian approach and reasoned as follows. In our data, each cell is a vector of counts, with one element for every gene. These counts can be viewed as a multinomial sample from some underlying distribution of gene expression. Since each state of the standard differentiation trajectory is defined by a particular gene expression distribution, we can ask: what is the probability a given direct programming protocol cell was sampled from each standard differentiation trajectory cluster? In this usage, probability amounts to a measure of similarity, with high probability indicating high similarity. We obtained similar results working in either a PV-gene expression space, or considering all genes expressed above a minimum counts threshold. The results of this analysis using PV-genes are presented in Figure 3D. We also obtained similar (but more noisy) results using cosine similarity as an alternative distance metric (not shown). The specific steps of our computation are as follows:

1. Extract a set of genes with which to make comparisons (either PV-genes or all genes above a minimum expression threshold).
2. For each cluster of the standard trajectory, calculate the probability of observing a given gene (i.e. the fraction of counts in that cluster from the gene). For genes that are not detected add 1e-07 total counts.
3. For each cell in the direct programming trajectory, calculate the log-likelihood that it was drawn from each of these clusters. This log-likelihood is from the multinomial distribution function using the probabilities obtained in step 2.
4. Identify and tally the maximum likelihood assignments of all direct programming cells. Normalize raw assignments so that they sum to 100 (giving the percentage). Plot the percentage of direct programming cells assigned to each standard protocol state.

### Cell cycle gene expression analysis

Cell-cycle activity can be estimated from a cell cycle associated gene expression signature; populations that express higher average levels of cell cycle genes are most likely cycling at higher frequency than a population with lower level expression of these genes. In Fig. 4b and Fig. S4 we performed an analysis in this spirit to determine which parts of the DP and SP differentiation trajectories appeared to be proliferative, and to estimate where cells are exiting the cell cycle. We computed a proliferation score that was the aggregate expression of a panel of 21 cell cycle related genes: Aurka, Top2a, Ccna2, Ccnd1, Ccnd2, Ccnd3, Ccne1, Ccne2, Ccnb1, Cdk4, Cdk6, Cdk2, Cdk1, Cdkn2b, Cdkn2a, Cdkn2c, Cdkn2d, Cdkn1a, Cdkn1b, Cdkn1c, Mcm6, Cdc20, Plk1, and Pcna. We also computed a cell cycle exit score on the basis of the aggregate expression of a panel of 4 tumor suppressor genes that inhibit the cell cycle: Cdkn1c, Cdkn1b, Cdkn1a, and Cdkn2d. In Fig. S4 we show the expression of representative individual genes from this score; in general cell cycle genes were correlated with each other in their expression over cells, as were cell cycle exit genes.

### Comparison of motor neurons *in vitro* with primary motor neurons using single-cell data

How does the transcriptional state of motor neurons produced by both protocols compare to that of motor neurons *in vivo*? To answer this question we leveraged the ability of single-cell RNA sequencing to compare cell states even within populations that are not pure (also see below for functional comparisons of the phenotypes). We performed three analyses. First, we computed the cosine similarity between the centroids of each cell state in both protocols and primary motor neurons (Fig. 5B). The specific steps of this analysis were as follows:

1. Combine all cell states from both protocols, and from HB9+ E13.5 primary tissue, extract PV-genes (as described above), and z-score normalize their expression.
2. Calculate the centroid of every cluster from each trajectory, and from primary motor neurons, in this PV-gene space.
3. Compute the cosine similarity for all *in vitro* populations versus primary motor neurons, and visualize as a bar graph.

Second, we performed a co-clustering analysis (Supp. Fig. 6). If cells cluster together this is an indication of similarity. An advantage of co-clustering is that it allows one to make statements at a single cell level about the fraction of cells within a state that resemble another state, for example. The specific steps of our coclustering analysis were:

1. For standard protocol, direct programming, and primary HB9+ cells, filter for neurons: in this analysis we defined neurons as cells with > 2.5 UMIs from universal neuronal marker Tubb3.
2. Perform tSNE and local density gradient clustering (as above; Supp. Fig. 6A).
3. For every condition, count the fraction of cells belonging to each cluster. Similar states should belong to the same cluster.
4. Visualize: we chose to use a pie chart.

Third, we performed differential gene expression analysis, comparing the most mature motor neuron states from each protocol with primary HB9+ motor neurons (Supp. Fig. 7). This analysis was performed as described above. We also performed differential expression analysis of these populations using bulk microarrays as validation (see below).

### qPCR

RNA was isolated using Qiagen Rneasy Plus Kit. Purified RNA was then reverse transcribed using Biorad’s iSCRIPT cDNA synthesis kit. Quantitative PCR was performed using Bio-rad’s SYBR green Supermix on a CFX96 Real-Time PCR system.

### Immunostaining

Antibodies used for immunostaining were anti-Tubb3 (Cell Signaling D71G9; 1:100), anti-Map2 (Sigma M9942; 1:200), anti-VACht (Synaptic Systems 139 103; 1:200), anti-Isl1 (Abcam ab109517; 1:1000), and anti-Hb9 (DSHB 81.5C10; 1:100). Differentiated cells were fixed in 4% paraformaldehyde for 20 minutes and then permeabilized with 0.1% Triton-X for 15 minutes. After primary incubation for 1 hour, samples were labeled with a secondary antibody conjugated to AlexaFluor647. Samples were co-stained with DAPI before imaging on a Nikon Eclipse TE2000-E microscope.

### Electrophysiology

All recordings were carried out at room temperature within 6 days of plating the neurons in 35 mm dish. Whole-cell voltage clamp recordings were made with a Multiclamp 700B amplifier (Molecular Devices) and patch pipettes with resistances of 2-3 ΜΩ. Pipette solution was 135 mM K-Gluconate, 10 mM KCl, 1 mM MgCl_2_, 5 mM EGTA, 10 mM HEPES, pH 7.2, adjusted with NaOH. The external solution was 140 mM NaCl, 5 mM KCl, 2 mM CaCl_2_, 2 mM MgCl_2_, 10 mM HEPES, and 10 mM D-glucose, pH 7.4, adjusted with NaOH. We used gravity perfusion system connected with Perfusion Pencil^®^ with Multi-Barrel Manifold Tip (AutoMate Scientific) to externally apply 0.5 μM tetrodotoxin, 100 μM AMPA, kainate, GABA, or glycine to the cells. Command protocols were generated and data was digitized with a Digidata 1440A A/D interface with pCLAMP10 software.

### Co-culture muscle contraction assays

C2C12 myoblasts were grown in 10%FBS+DMEM media and then differentiated into myotubes by incubating in differentiation medium (2% horse serum + DMEM). After myotubes were formed, the neurons were dissociated by trypsinization and reseeded on top of the differentiated muscle to allow contractions to develop. Video of contractions were taken using Metamorph software and manually counted over 5 second intervals. For stopping assay, 300 μM Tubocurarine (Sigma) was added to the media as an acetylcholine competitor. For labeling of acetylcholine receptors, bungarotoxin (Invitrogen) was used after cells were fixed with paraformaldehyde. Similarly to C2C12 cells, ES cells over-expressing MyoD were also differentiated to myotubes using differentiation medium and subjected to co-culture with neurons.

### Microarray Analysis

The mRNA of undifferentiated iNIL ES cells and NIH 3T3 cells (grow in DMEM+10%FBS) were collected and purified by RNA extraction using RNeasy Plus Extraction Kit (Qiagen). Neurons differentiated by either protocol were first sorted by flow cytometry on a BD FACSaria II machine (Beckton Dickinson, USA) to collect Hb9::GFP+ cells, and then were subjected to RNA extraction in a similar fashion. Collected RNA was then amplified and hybridized to Affymetrix GeneChip Mouse Transcriptome Arrays (MTA 1.0). Results were processed by the Children’s Hospital Microarray Core Facility, and were analyzed using Affrymetrix’s Transcriptome Analysis Console and Expression Console software.

